# Distinct System-Level Computations Underlie Perceptual Variation Across the Visual Field

**DOI:** 10.1101/2025.09.19.677418

**Authors:** Shutian Xue, Antoine Barbot, Jared Abrams, Qingyuan Chen, Marisa Carrasco

**Author notes:** Correspondence: Shutian Xue, Marisa Carrasco. These authors contributed equally to this manuscript (Shutian Xue, Antoine Barbot).

## Abstract

Human visual perception for basic dimensions varies with eccentricity and polar angle, influencing daily activities such as reading, searching and scene perception. We investigated whether and how system-level computations that transform visual input into perception underlie these heterogeneities. Using the equivalent noise method and perceptual template model, we estimated gain, internal noise, and nonlinearity for orientation discrimination across eccentricity (fovea, parafovea and perifovea) and around polar angle. Participants discriminated the orientation of Gabors embedded in dynamic white noise and showed the expected variations across eccentricity and around polar angle. Importantly, visual performance declined with eccentricity due to decreased gain and nonlinearity and increased internal noise. Observers with stronger eccentricity effects showed greater gain decrease. Only gain varied with polar angle—higher along the horizontal than vertical meridian, and lower than upper vertical meridian—paralleling performance asymmetries. This dissociation aligns with known variations in neuronal count and tuning, suggesting that neural correlations and neural noise contribute to these system-level computations. By revealing distinct system-level computations underlying the eccentricity effect and polar angle asymmetries, our findings link perceptual heterogeneity across the visual field and neural architecture and provide insights into how the human brain encodes information under neural constraints.

**Significance Statement:** Human visual performance varies across eccentricity–distance from gaze–and around polar angle –circular dimension. Retinal factors and cortical surface area account for eccentricity, but only partially for polar angle variations. Here, we show that system-level computations—how signal are amplified and how noisy the system is—distinctly underlie these perceptual variations. Decline across eccentricity stems from both reduced gain and nonlinearity and elevated internal noise, whereas variation around polar angle arises solely from gain difference. This dissociation implies fundamental differences in information processing along two axes of the visual field. Our results provide a computational link between behavior and its neural bases and highlight the importance of considering both eccentricity and polar angle in modeling and understanding human perception.

## Introduction

Visual performance declines with eccentricity^1–3^ and varies systematically around polar angle at isoeccentric locations. Performance is superior along the horizontal than vertical meridian— horizontal-vertical anisotropy (HVA)—and along the lower than upper vertical meridian—vertical meridian asymmetry (VMA) (**Fig. 1A**; review^3^). These polar angle asymmetries are ubiquitous across multiple visual functions, from early vision, such as contrast sensitivity (e.g^4–8^) and acuity (e.g^8,9^), to mid-level vision, such as texture segmentation^10^ and crowding^11^, and high-level vision, such as visual short-term memory^12^ and face perception^13^. Their magnitude is equivalent to doubling^14,15^ or tripling^7^ eccentricity, and they persist under both binocular or monocular viewing^9,14^. These variations were amplified at more peripheral locations^7,14,16–18^ increase at higher spatial frequencies^4,14,15^, and remain robust even in the presence of attention^4,10,14,19,20^. Here, we investigate the computations by which the visual system gives that to this pervasive and resilient pattern of performance across the visual field.

**Figure 1.**
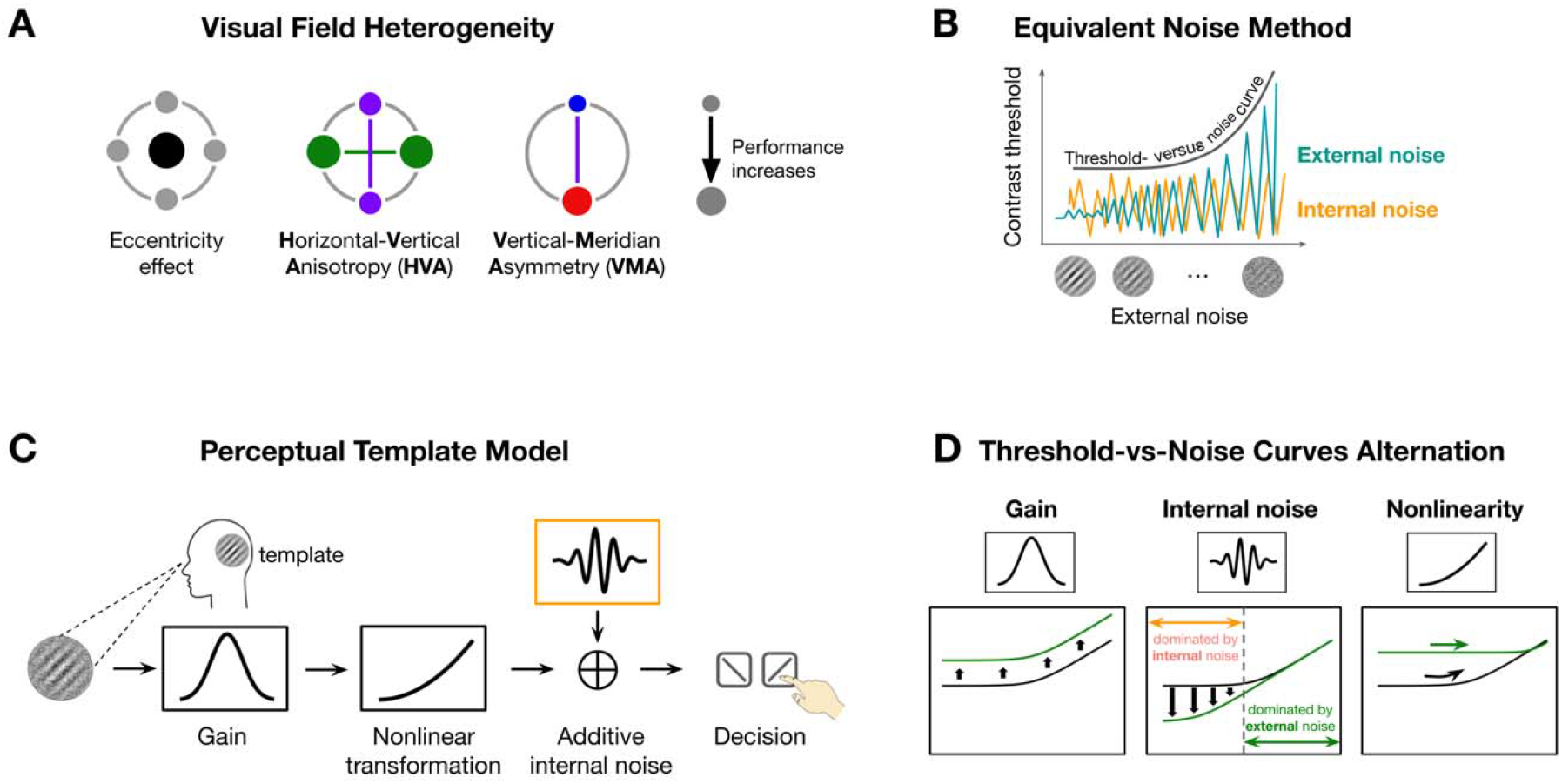
Schematics of key concepts. **(A)** A schematic of visual field heterogeneity in performance. The size of the dot indicates performance (bigger dots = better performance). From left to right we illustrate the eccentricity effect (fovea is better than farther eccentricities), horizontal-vertical anisotropy (HVA, the horizontal is better than the vertical meridian), and vertical-meridian asymmetry (VMA, the lower is better than the upper vertical meridian). **(B)** The equivalent noise method infers neural computations by mapping performance (e.g., contrast threshold) as a function of external noise and observing how performance is altered by both external (dark green) and internal (orange) noise. The performance decrease with higher external noise levels is captured by the threshold-vs-noise (TvN) curve. **(C)** The Perceptual Template Model (PTM) includes three system-level computations—gain, additive internal noise and nonlinearity. A noisy stimulus is filtered by a perceptual template, and its contrast is scaled by a gain parameter, transformed nonlinearly, and corrupted by internal noise before decision-making. **(D)** Proposed computations influence performance in distinct ways. Differences in gain scale the entire curve without altering its shape, reflecting proportional changes in how the system filters both signal and noise; Changes in internal noise primarily affect thresholds at low external noise levels, where internal variability dominates. As external noise increases, performance converges across locations. Nonlinearity determines the curvature of the function: higher nonlinearity results in a more curved (nonlinear) function.

Both the eccentricity effect and polar angle asymmetries are related to better optical quality^21^, higher density of cones^22,23^ and midget retinal ganglion cells^24,25^ and larger surface area in primary visual cortex (V1) (eccentricity^26–31^; HVA/VMA^28–30,32^). Importantly, V1 surface area correlates with contrast sensitivity around polar angle^32^, indicating a link between cortical anatomy and visual perception at the level of individual observer.

However, retinal and cortical factors are insufficient to account for polar angle asymmetries. An encoding model simulating orientation discrimination based on spatial variations in both optical and retinal factors explained only 10% of VMA and 40% of HVA performance^33,34^. Furthermore, M-scaling the stimulus size as per a well-established cortical magnification factor eliminates contrast sensitivity differences across eccentricity (e.g^7,35^), but not around polar angle^7^.

Here, we investigated whether system-level computations reflecting the pooled activity of neuronal populations activated by the stimulated visual field play a critical role. We examined three computations that limit visual performance in a noisy environment: gain —the system’s ability to amplify the signal^36–38^ (or modeled as efficiency^39^), nonlinearity—how the system transforms inputs in a nonlinear fashion^40,41^, and internal noise—the variability intrinsic to the visual system, independent of external stimulation^40–42^. Both gain and nonlinearity are template properties reflecting the system’s ability to extract task-relevant information from stimuli. We used the equivalent noise method to probe these system-level computations by introducing external noise (e.g^40–45^) and infer the three computations by observing how performance changes (**Fig. 1B**). We constructed threshold-versus-noise curves at different locations to capture how performance is jointly influenced by external noise and system-level computations.

The perceptual template model quantifies the three system-level computations (**Fig. 1C**). We used a simplified version of the original model developed by Lu and Dosher (e.g^40,41,43^), which retains its explanatory power. First, the noisy stimulus is filtered through a perceptual template tuned to the signal’s orientation and spatial frequency (SF). This step is scaled by the gain parameter, such that better template-signal alignment yields higher gain. Then, the template-matched output is passed through a nonlinear transformation, typically a power function. Next, this transformed signal is corrupted by additive internal noise. Finally, a decision is made based on the noisy signal.

These neural computations have been linked to neurophysiological properties. Higher gain reflects stronger template alignment, which may depend on smaller receptive fields and sharper SF tuning^46^, thereby supporting higher sensitivity. Neurons have intrinsic variability due to factors such as fluctuations in firing rate and membrane potential^47^. At the system level, internal noise is shaped by neural correlations: when neural activity is more correlated among neurons, their noise is less likely to cancel out through averaging, leading to greater internal noise at the system level^48,49^. Nonlinearity is shaped by both retinal and cortical processes, including synaptic rectification^50^ and divisive normalization^51^.

Each computation influences performance in distinct ways (**Fig. 1D**). When gain differs between two locations, the system’s ability to filter both signal and noise changes proportionally, resulting in a vertical shift of the threshold-versus-noise curve without altering its shape. Differences in internal noise impact performance the most at lower external noise levels, where internal noise dominates. As external noise becomes more dominant, performance at higher external noise becomes indistinguishable across locations. Finally, nonlinearity determines the curvature of the function.

To investigate how these system-level computations shape visual field heterogeneity, we assessed whether they vary throughout the visual field. Previous studies have shown that internal noise increases across eccentricity^52–54^. Neuroimaging studies have shown that the size of surface area decreases with eccentricity (e.g^26,27,31^). These studies averaged data around polar angle; thus, it is unknown whether polar angle modulates these computations. Based on the anatomical evidence that the decline of cone and retinal ganglion cell density^22–25^ and cortical surface area^28^ across eccentricity is steeper along the vertical than horizontal meridian, as well as behavioral findings that the eccentricity effect is more prominent along the vertical than horizontal meridian (e.g^5,16,55,56^) and a robust HVA (e.g^6,8^), we hypothesize that the eccentricity effect as well as the gain and internal noise underlying this effect will vary more along the vertical than horizontal meridian.

It is unknown whether gain and internal noise also vary around polar angle. Anatomical and neuroimaging measurements, including cone and retinal ganglion density^22,23^, as well as V1 cortical surface area^28–32^, population receptive field size^28,31^, and BOLD signal amplitude^11,57^, contribute to the behavioral polar angle asymmetries^33,34^. Accordingly, both gain and internal noise may vary around polar angle. However, magnifying stimulus size only reduced variation around polar angle, suggesting that differences in V1 cortical surface area do not fully explain polar angle asymmetries^7^. Here, we explored whether in addition to retinal and cortical differences, polar angle asymmetries also reflect differences in gain and internal noise, or whether the underlying computations differ from those underlying the eccentricity effect. We also assessed nonlinearity across eccentricity and around polar angle, as it reflects how the visual system amplifies or compresses neural responses and plays a critical role in shaping the threshold-versus-noise curve.

Second, we examined whether variations in gain, internal noise, and nonlinearity across locations contribute to performance heterogeneity. Specifically, we asked whether the magnitude of variation in system-level computations—quantified by a location difference index—predicts corresponding differences in performance. By quantifying the impact of each computation on performance^54^, this analysis provides further evidence that system-level computations play a mechanistic role in shaping visual perception throughout the visual field.

## Results

Observers performed an orientation discrimination task at nine visual field locations: fovea, four cardinal meridians (left and right horizontal meridian and upper and lower vertical meridian) at 4° and 8° eccentricity (**Fig. 2A**). The target signal was a tilted Gabor embedded in dynamic white noise with a SF of either 4 or 6 cpd. We varied the contrast of Gabor and noise across trials to map contrast threshold as a function of external noise, location and SF (**Fig. 2B**). We estimated three parameters—gain, additive internal noise, and nonlinearity—by fitting the perceptual template model to threshold-versus-noise curves at each location (**Fig. 2C**). Finally, we assessed how each parameter varies across eccentricity and around polar angle, and contributes to location differences of performance.

**Figure 2.**
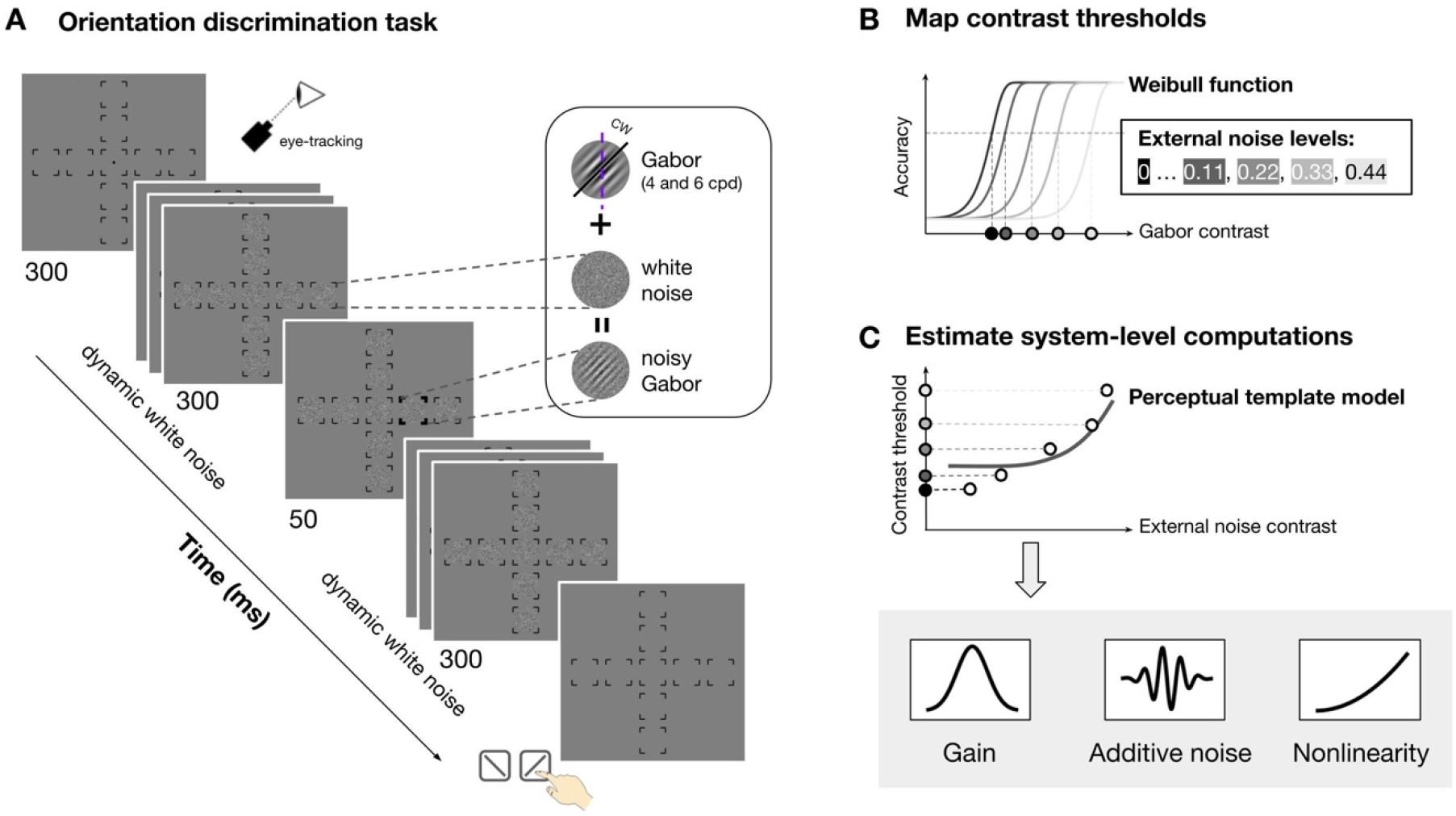
Stimulus, procedure, and model fitting. **(A)** Orientation discrimination task. Observers saw a streak of dynamic white noise presented at nine locations (fovea, four polar angle at 4° and 8° eccentricity) both before and after seeing a Gabor signal presented at one location. They reported whether the Gabor was tilted clockwise or counterclockwise and received feedback. The signal location, noise contrast and signal contrast varied across trials. The Gabor SF varied across sessions (4 and 6 cpd, counterbalanced). Eye fixation was enforced by an eye tracker. **(B)** To derive contrast thresholds at each location and each noise level, we fitted a Weibull function to accuracy as a function of Gabor contrast (binned into 10 bins) for each noise level (indicated by lines of different shades) and estimated threshold (indicated by the dots of different shades at x-axis). **(C)** We fitted the perceptual template model to the thresholds across external noise to derive the three computations: gain, additive internal noise and nonlinearity. The solid dots indicate the threshold estimated in **(B)**.

### Visual field heterogeneities in the absence of external noise

We first verified the visual field heterogeneity—the eccentricity effect and polar angle asymmetries—in absence of external noise and examined how SF modulated their magnitudes. **Fig. 3** shows threshold-versus-noise curves for each location group at 4 and 6 cpd. Only thresholds estimated at 75% accuracy are shown, for illustration purposes. We focused on thresholds estimated at external noise equal to 0, referred to as *performance*. We fit linear mixed-effects models to performance and conducted Type III ANOVAs to examine interactions between location (eccentricity/meridian/specific location) and SF (4 and 6 cpd) and their main effects (Materials and Methods, equation (9)).

**Figure 3.**
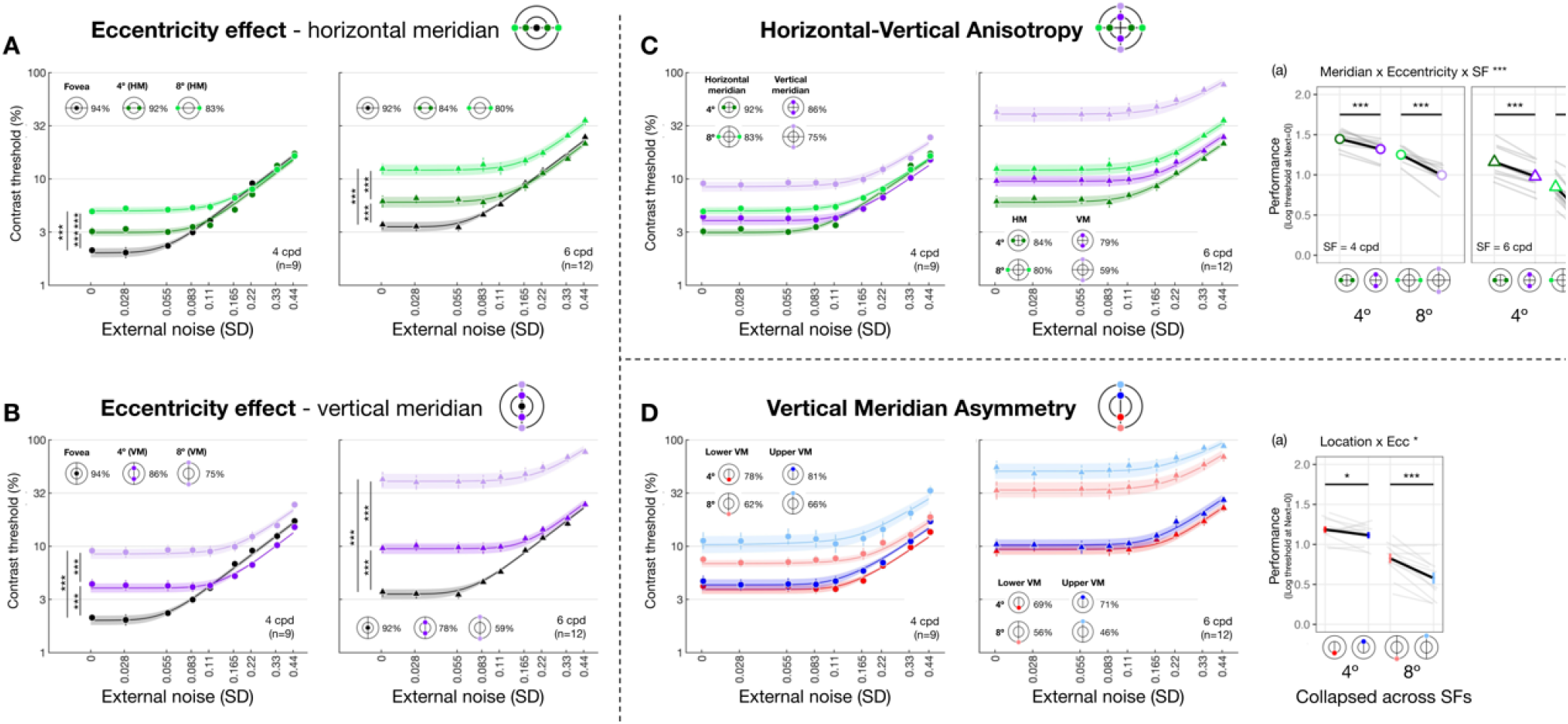
Threshold-vs-noise curves and performance throughout the visual field. Each panel plots thresh-vs-noise curve for each location group to visualize one of the three visual field heterogeneities: eccentricity effect along the horizontal **(A)** and vertical meridian **(B)**, HVA **(C)** and VMA **(D)**. Dots and error bars represent group-averaged contrast thresholds (±1 SEM) across nine levels of external noise. Many error bars are smaller than the symbols. Shaded bands are PTM model predictions (±1 SEM). Colors denote visual field locations, indicated by icons in each panel, and corresponding R^2 values for PTM fitting are shown alongside. Dot shapes indicate SF (circles=4 cpd, triangles=6 cpd), plotted separately in the left and right panels within each dashed cell. **(A, B)** Threshold-vs-noise curves at the fovea, and at 4° and 8° eccentricity along the horizontal **(A)** and vertical **(B)** meridian. To examine eccentricity effects, we compared *performance*—defined as the absolute value of the log contrast threshold at zero external noise— between each pair of eccentricities (fovea vs. 4°, fovea vs. 8°, and 4° vs. 8°). Vertical comparison lines indicate the pairs tested; asterisks denote Bonferroni-corrected *p*-values (\*\*\**p* < .001, \*\**p* < .01, \**p* < .05, ns, *p*>.1). **(C)** Threshold-vs-noise curves at the horizontal and vertical meridians at both 4° and 8° eccentricity. Subpanel (a) plots performance as a function of meridian × eccentricity, separately for 4 and 6 cpd, illustrating how both eccentricity and SF modulate the HVA. Group means (±1 SEM) are plotted as colored dots; individual subjects’ performance (median of 1,000 bootstraps) is shown in grey lines. We focused on comparisons between horizontal and vertical meridian, indicated by horizontal comparison lines. Asterisks denote significance levels from post-hoc pairwise tests. **(D)** Threshold-vs-noise curves at the lower and upper vertical meridians at 4° and 8° eccentricity. Subpanel **(a)** shows performance collapsed across SF as a function of meridian × eccentricity, revealing eccentricity-dependent modulation of the VMA. Formatting follows that of **(C)**.

We observed an interaction between eccentricity and SF along both the horizontal meridian (*F*(2, 45.7) = 7.277 [1.623, 9.959], *p*_*obs*_ = 0.002, 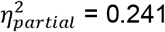 [0.067, 0.303]; **Fig. 3A**) and the vertical meridian (*F*(2, 45.7) = 29.089 [10.615, 30.430], *p*_*obs*_ < 0.001, 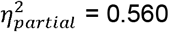 [0.318, 0.570]; **Fig. 3B**). Post-hoc pairwise comparisons revealed that performance decreased with eccentricity along both meridians and both SFs (all eccentricity pairs had *p*_*emp*_ < 0.001). This decrement was more pronounced for 6 than 4 cpd and for vertical than horizontal meridian (averaged Cohen’s *d*: Vertical-6 cpd > Vertical-4 cpd > Horizontal-6 cpd > Horizontal-4 cpd). These results confirm the well-established eccentricity effect, in which a decline in performance is more pronounced at higher SFs and along the vertical than the horizontal meridian.

We then compared performance between the horizontal and vertical meridian (**Fig. 3C-a**) and examined the modulation of eccentricity (4° and 8°) and SF (4 and 6 cpd) (Materials and Methods, equation (10)). We found a three-way interaction among meridian, eccentricity and SF (*F*(1,64.7) = 6.728 [0.178, 10.399], *p*_*obs*_ = 0.012, 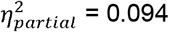 [0.003, 0.139]). Post-hoc analyses revealed that performance was higher at the horizontal than the vertical meridian for all four SF x eccentricity combinations, confirming a HVA in performance. The largest HVA was observed at 6 cpd x 8° eccentricity (*t*(65.0) = 14.030 [7.869, 14.100], *p*_*emp*_ < 0.001, Cohen’s *d* = 17.702), followed by 4 cpd x 8° (4 cpd; *t*(65.0) = 5.890 [3.060, 7.730], *p*_*emp*_ < 0.001, Cohen’s *d* = 8.122). The asymmetries at 4° eccentricity were smaller but still statistically significant for both 4 cpd (*t*(65.0) = 3.010 [1.410, 10.399], *p*_*emp*_ < 0.001, Cohen’s *d* = 5.907) and 6 cpd (*t*(65.0) = 5.090 [3.060, 5.390], *p*_*emp*_ < 0.001, Cohen’s *d* = 5.225). In sum, the performance HVA was robust and stronger at further eccentricity and higher SF.

Along the vertical meridian (**Fig. 3D-a**), the three-way interaction (location x eccentricity x SF) was not significant (*p*_*obs*_ > 0.1). Location interacted with eccentricity (*F*(1,64.6) = 4.400 [0.945, 9.634], *p*_*obs*_ = 0.040, 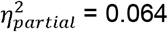 [0.014, 0.130]) but not with SF (*p*_*obs*_ > 0.1), indicating that performance variation along the vertical meridian was modulated by eccentricity, but not by SF. Post-hoc analysis showed that performance was better at lower than upper vertical meridian at both 4° (*t*(65.0) = 1.300 [0.139, 1.890], *p*_*emp*_ = 0.044, Cohen’s *d* = 2.258) and 8° eccentricity (*t*(65.0) = 4.270 [2.469, 5.060], *p*_*emp*_ < 0.001, Cohen’s *d* = 5.806), confirming a VMA in performance.

Together, we observed robust eccentricity effects and polar angle asymmetries in performance. Effects are more pronounced at the higher SF and farther eccentricities.

### Gain, additive noise and nonlinearity vary throughout the visual field

To examine how computations underlie the observed visual field heterogeneity, we first assessed whether and how each parameter varied throughout the visual field. We fitted linear mixed-effects models (Materials and Methods, equation (11-12)) to each parameter and conducted Type III ANOVAs to test interactions between locations and SF and their main effects. Matching between a parameter’s variation pattern with typical visual field performance patterns—eccentricity effect, HVA and VMA—suggests that the corresponding computation underlies these visual field heterogeneities.

#### Gain, additive noise and nonlinearity vary across eccentricity

When eccentricity increases from fovea to 4°, at 4 cpd the curve differs at lower external noise levels but converges at higher levels (**Fig. 3A**), indicating increased internal noise at parafoveal locations (**Fig. 1D**, *middle*). As eccentricity increases further to 8° eccentricity, performance decreased across all levels of external noise, suggesting a reduction in gain at peripheral locations (**Fig. 1D**, *left*). Based on these qualitative differences, we quantitatively compared gain and internal noise across the visual field. For gain, along the horizontal meridian (**Fig. 4A**), eccentricity interacted with SF (*F*(2, 57.0) = 6.032 [0.509, 9.016], *p*_*obs*_ = 0.004, 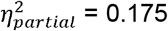 [0.018, 0.264]). Post-hoc pairwise comparisons showed that for 6 cpd, 8° eccentricity had lower gain than both fovea (*t*(46.7) = 3.220 [0.799, 3.830], *p*_*emp*_ = 0.048, Cohen’s *d* = 3.248) and 4° eccentricity (*t*(46.7) = 4.680 [1.430, 4.620], *p*_*emp*_ = 0.036, Cohen’s *d* = 4.400); but for 4 cpd, gain was similar across eccentricity. For additive internal noise (**Fig. 4B**), there was a main effect of eccentricity (*F*(2, 45.1) = 34.842 [10.783, 38.066], *p*_*obs*_ < 0.001, 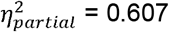 [0.307, 0.623]): fovea was lower than 4° (*t*(46.3) = −5.310 [−5.991, −2.350], *p*_*emp*_ = 0.006, Cohen’s *d* = −4.380), which in turn was lower than 8° eccentricity (*t*(46.3) = −2.920 [−3.540, −1.110], *p*_*emp*_ < 0.001, Cohen’s *d* = −3.611). Lastly, nonlinearity (**Fig. 4C**) also varied with eccentricity along the horizontal meridian (*F*(2, 46.6) = 11.916 [1.653, 14.295], *p*_*obs*_ < 0.001, 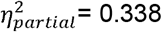 [0.065, 0.379]), being higher at the fovea than 8° eccentricity (*t*(46.3) = 4.870 [1.640, 5.221], *p*_*emp*_ = 0.006, Cohen’s *d* = 3.523).

**Figure 4.**
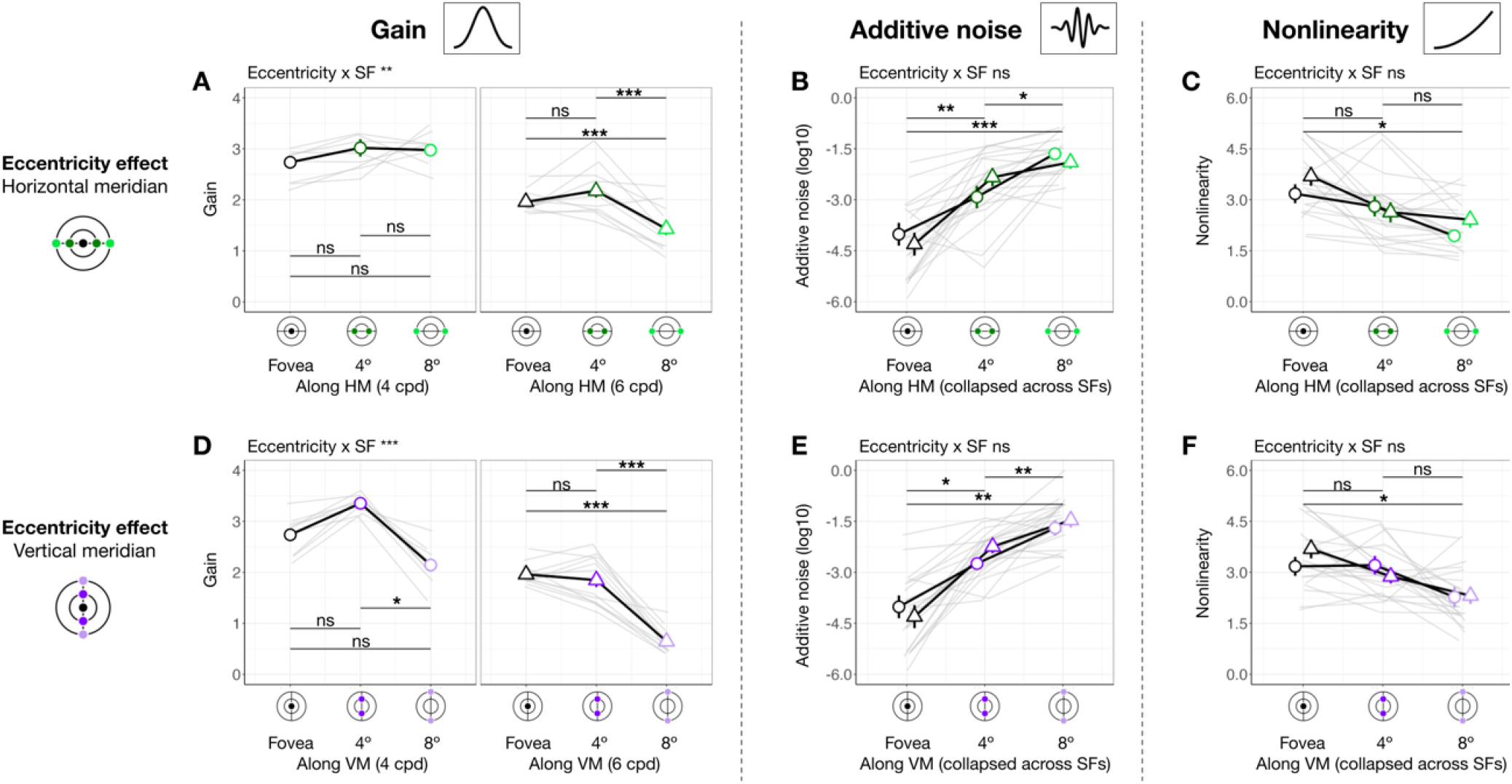
Gain, additive noise and nonlinearity vary across eccentricity. Each panel shows group means (±1 SEM) of estimated model parameters—gain **(A, D)**, additive noise **(B, E)**, and nonlinearity **(C, F)**—as a function of eccentricity along the horizontal (top row) and vertical (bottom row) meridian. Grey lines represent individual subject data (median of 1,000 bootstraps). Asterisks above horizontal comparison lines indicate Bonferroni-corrected post-hoc significance levels (***p < .001, *p < .05, ns p > .1). Parameters that showed a significant interaction between eccentricity and SF (e.g., gain in panels **A** and **D**) are plotted separately for 4 and 6 cpd. Parameters showing only a main effect of eccentricity without such interaction are collapsed across SFs (additive noise in panels B and E, and nonlinearity in panels **C** and **F**).

A similar pattern emerged at the vertical meridian (**Fig. 4D-F**). For gain, eccentricity interacted with SF (*F(*2, 57.0) = 11.098 [0.657, 10.210], *p*_*obs*_ < 0.001, 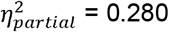 [0.028, 0.297]; **Fig. 4D)**. At 6 cpd, Gain was lowest at 8° eccentricity and lower than at 4° (*t*(46.7) = 8.720 [3.229, 8.011], *p*_*emp*_ < 0.001, Cohen’s *d* = 7.123) and at the fovea (*t*(46.7) = 9.870 [3.449, 8.752], *p*_*emp*_ < 0.001, Cohen’s *d* = 8.542). Moreover, at 4 cpd, gain was lower at 8° than 4° eccentricity (*t*(46.7) = 5.990 [1.339, 7.720], *p*_*emp*_ = 0.024, Cohen’s *d* = 3.418). Additive noise increased with eccentricity (**Fig. 4E**), (*F*(2, 45.3) = 48.393 [13.872, 46.819], *p*_*obs*_ < 0.001, 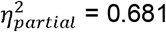 [0.361, 0.673]) without interacting with SF (*p*_*obs*_ > 0.1). Additive noise was lower at the fovea than at 4° (*t*(46.2) = −6.130 [−6.780, −2.569], *p*_*emp*_ = 0.024, Cohen’s *d* = −4.343), which in turn was lower than at 8° eccentricity (*t*(46.2) = −3.590 [−4.120, −1.000], *p*_*emp*_ = 0.006, Cohen’s *d* = −3.215). Similarly, nonlinearity varied across eccentricity (*F*(2, 47.0) = 6.787 [0.745, 11.888], *p*_*obs*_ = 0.023, 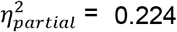 [0.030, 0.337]; **Fig. 4F**), being higher at the fovea than 8° eccentricity (*t*(46.4) = 3.600 [0.930, 4.580], *p*_*emp*_ = 0.024, Cohen’s *d* = 2.883). Overall, the effect size of gain, but not of internal noise or nonlinearity, was higher along the vertical than the horizontal meridian, indicating a more pronounced gain decline across eccentricity along the vertical meridian.

In sum, gain decreased with eccentricity from fovea to 8°, and more so from 4° to 8°—with this effect being the most pronounced along the vertical meridian and for 6 cpd. Additive noise increased with eccentricity, and nonlinearity was higher at fovea than 8° eccentricity, similarly along both meridians.

#### Gain, but not internal noise or nonlinearity, varies around polar angle

We next assessed whether gain differed between horizontal and vertical meridian and whether this was modulated by eccentricity and SF (**Fig. 5A**). There was no three-way interaction (*p*_*obs*_ > 0.1), but meridian interacted with eccentricity (*F*(1, 56.9) = 13.960 [1.503, 25.598], *p*_*obs*_ < 0.001, 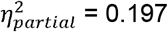 [0.022, 0.290]; **Fig. 5A-a**) and SF (*F*(1, 56.9) = 4.430 [0.006, 8.976], *p*_*obs*_ = 0.040, 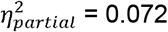 [0, 0.123] **Fig. 5A-b**). Post-hoc analyses showed that gain was higher at the horizontal than the vertical meridian at 8° eccentricity (*t*(65.5) = 6.130 [1.870, 6.560], *p*_*emp*_ < 0.001, Cohen’s *d* = 4.297) but not at 4° (*p*_*emp*_ > 0.1), consistent with the HVA pattern for performance. Similarly, gain reflected the HVA pattern at 6 cpd (*t*(65.5) = 5.370 [1.560, 4.650], *p*_*emp*_ = 0.008, Cohen’s *d* = 4.649), but not at 4 cpd (*p*_*emp*_ > 0.1). In sum, gain showed a pattern consistent with HVA, with stronger effects at greater eccentricities and higher SF. Neither additive noise (**Fig. 5B**) nor nonlinearity (**Fig. 5C**) differed between horizontal and vertical meridian (*p*_*obs*_ > 0.1) at any eccentricity or SF. Note that the lower additive noise at 4° than 8° eccentricity (**Fig. 5B**) was already captured by the eccentricity difference (**Fig. 4B**). These findings suggest that HVA arises specifically from variation in gain.

**Figure 5.**
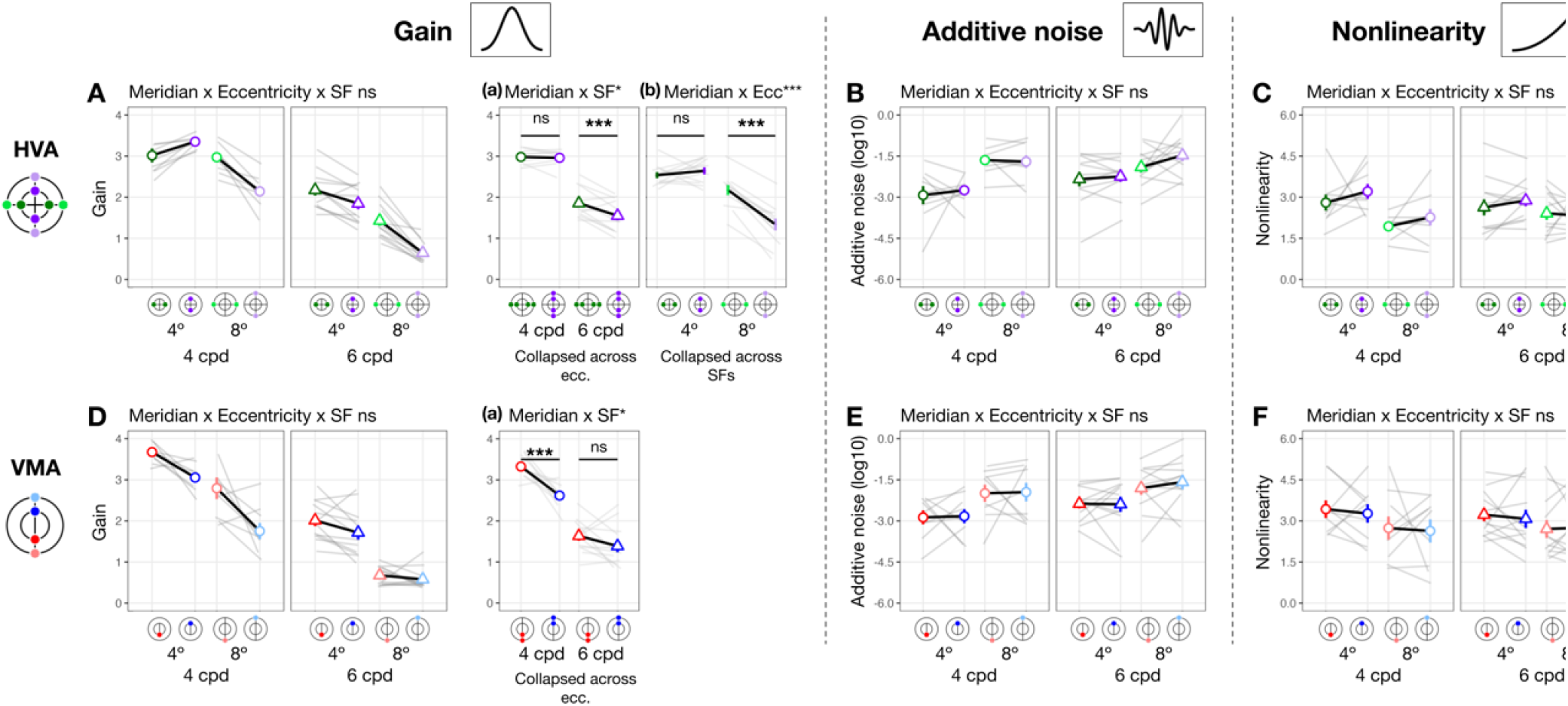
Gain **(A and D)**, but not internal noise **(B and E)** or nonlinearity **(C and F)**, varies around polar angle. We plot estimated parameters as a function of location × eccentricity pairs, separately for 4 and 6 cpd. To highlight post hoc comparisons from significant two-way interactions for gain **(A and D)**, we collapsed across SF to illustrate the modulation effect by eccentricity (panels **A-a** and **B-a**), given a significant meridian/location × eccentricity interaction. Similarly, we collapsed across eccentricity to illustrate the modulation effect by SF (panels **A-b** and **B-b**), given a significant meridian/location × SF interaction. Figure format follows that of **Fig. 4**.

We next examined whether gain differed between the lower and upper vertical meridian (**Fig. 5D)**. Location interacted with SF (*F*(1, 76.0) = 6.469 [0.071, 12.967], *p*_*obs*_ = 0.013, 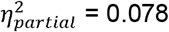 [0.001, 0.165] **Fig. 5D-a**), and gain was higher at lower than upper vertical meridian at 4 cpd (*t*(65.5) = 4.410 [1.280, 5.580], *p*_*emp*_ < 0.001, Cohen’s *d* = 3.374), but not at 6 cpd (*p*_*emp*_ > 0.1). This result means that VMA for gain was stronger at 4 than 6 cpd. The lack of VMA at a higher SF is because for some observers, gain at 6 cpd (particularly at 8° eccentricity) was near the floor level (close to 1), limiting any differences. There was no three-way interaction or interaction between location and eccentricity (*p*_*obs*_ > 0.1). Neither additive noise (**Fig. 5E**) nor nonlinearity (**Fig. 5F**) differed between the upper and lower vertical meridian (*p*_*obs*_ > 0.1). Together, these findings indicate that differences in gain, rather than in internal noise or nonlinearity, may underlie VMA.

In sum, gain alone consistently varies around polar angle, indicating that it is related to both HVA and VMA. These location-based effects are more pronounced at higher SFs. Distinct from the eccentricity effect, additive noise and nonlinearity did not vary around polar angle.

### Contribution: Gain, but not additive noise or nonlinearity, contributes to eccentricity effect

We examined the extent to which each system-level parameter—gain, additive internal noise, and nonlinearity—contributed to visual field asymmetries. First, we identified the direction in which each parameter modulated performance. Using linear mixed-effects models (Materials and Methods, equation (13)), we assessed the relation between performance and each parameter across seven locations in the visual field: the fovea, the horizontal meridian and the upper and lower vertical meridians at both 4° and 8° eccentricities.

As shown in **Fig. 6A-C**, locations with higher performance were associated with higher gain (slope = 0.090 [0.047, 0.130], *p*_*emp*_ < 0.001, conditional R-squared=0.75 [0.67, 0.80]; **Fig. 6A**) and lower additive noise (slope = −0.029 [−.042, −0.008], *p*_*emp*_ < 0.05, conditional R-squared=0.77 [0.70, 0.81]; **Fig. 6B**). Nonlinearity showed no significant relation with performance (slope = −0.019 [−0.043, 0.02], *p*_*emp*_ = 0.082, conditional R-squared = 0.79 [0.72, 0.82]; **Fig. 6C**). These findings confirm our hypothesis that stronger gain and lower internal noise improve performance, whereas nonlinearity has no effect.

**Figure 6.**
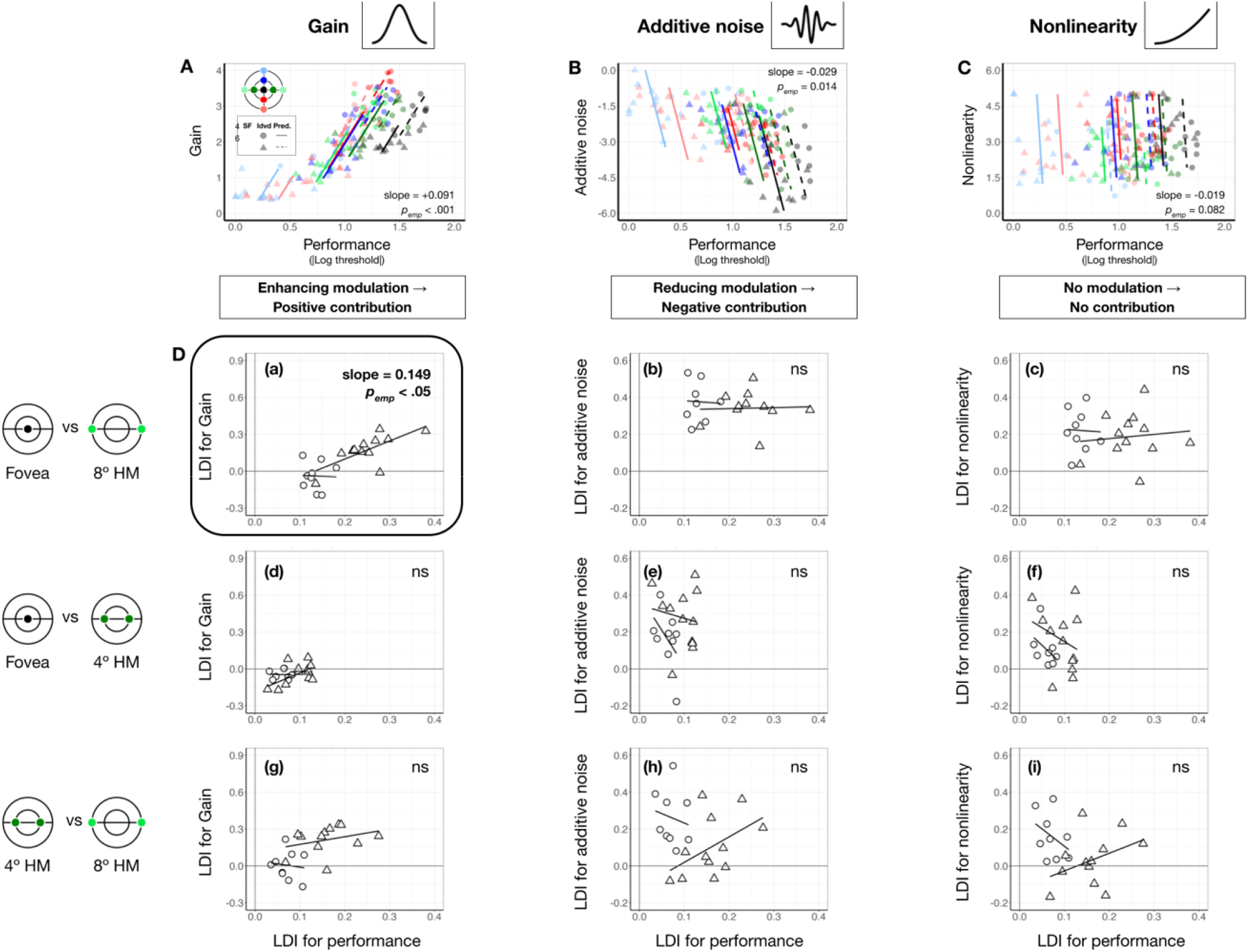
Gain contributes to performance differences between the fovea and 8° eccentricity. **(A-C)** Each panel shows estimated gain **(A)**, additive internal noise **(B)**, and nonlinearity **(C)** as a function of performance across locations and SFs. Each data point represents an individual location × SF combination (color-coded by location, shape-coded by SF). Colored regression lines show predictions from linear mixed-effects models. Medians of bootstrapped slopes and empirical *p*-values (*p*_*emp*_) are reported in the figure. Performance improves with increasing gain **(A)** and decreasing internal noise **(B)**, while nonlinearity shows no relationship with performance **(C). (D)** Quantify the contribution of each computation. Each panel plots the location difference index (LDI) of a parameter against the LDI of performance. Shapes denote individual observer data at 4 cpd (circles) and 6 cpd (triangles). Regression lines reflect linear mixed-effects model predictions for each SF. Median of bootstrapped regression slopes and empirical one-tailed *p*-values are shown in each panel. Subpanels **(a-c)** show gain results across three eccentricity pairs (fovea vs. 8°, fovea vs. 4° and 4° vs. 8°) along the horizontal meridian. A significant positive correlation appears only for the fovea vs. 8° pair **(D-a)**, especially at 6 cpd, indicating that gain differences contribute to the eccentricity effect. Subpanels **(d-f)** show additive internal noise, which shows no significant correlations with performance for any eccentricity pair. Subpanels **(g-i)** shows nonlinearity, which similarly shows no significant correlation with performance across eccentricity pairs.

The direction of each modulation informs the directionality of parameter contribution to location-based performance differences: Parameters that enhance performance should vary across locations in line with performance variation, whereas parameters that impair performance should vary in the opposite direction. We examined whether and how these location differences of each parameter are proportional to the heterogeneity in the expected direction (equation (14)).

Location variation was indexed by the difference between two locations divided by their sum, providing a normalized measure. In **Fig. 6D**, we plot location difference indices for each parameter against those for performance.

For gain, the only significant correlation was between the fovea and 8° along the horizontal meridian (slope = 0.164 [0.020,0.412], *p*_*emp, right-tailed*_ = 0.038, conditional R-squared = 0.75 [0.48, 0.93], **Fig. 6D-*a***). The effect was driven by the positive correlation at 6 cpd (*p*_*emp, right-tailed*_ = 0.030). No correlations were observed for the other eccentricity pairs (**Fig. 6D-*b* and *c*)**, or for internal noise and nonlinearity (all *p*_*emp*_ > 0.1, **Fig. 6D-*e-i***), despite their variations across eccentricities. These results suggest that gain, particularly at higher SF, is the primary factor underlying the eccentricity effect along the horizontal meridian.

In contrast, no correlation with polar angle differences was significant (*p*s_*emp*_>.1), meaning that although gain varies around polar angle, it does not contribute to polar angle asymmetries as much as to the eccentricity effect.

## Discussion

Using the equivalent noise method and perceptual template modeling, we mapped contrast sensitivity across the visual field and identified distinct computations that underlie performance variations across eccentricity –which have been examined primarily along the horizontal meridian or collapsed across polar angle (review^3^)– and around polar angle. With increasing eccentricity, all three computations varied: gain and nonlinearity decreased, whereas internal noise increased. Gain variations across eccentricity were stronger along the vertical than horizontal meridian, consistent with visual performance (e.g^5,16,55,58^), and they were also stronger at higher SFs, in line with previous studies^5,59,60^. Moreover, gain predicted contrast sensitivity differences between fovea and 8°, indicating that gain is a key driver of the eccentricity effect. In contrast, around polar angle, neither internal noise nor nonlinearity varied, only gain did—being higher along the horizontal than the vertical meridian and at the lower than the upper vertical meridian—mirroring the HVA and VMA in performance. Together, these findings indicate that polar angle asymmetries arise from a distinct computational profile.

We found that the eccentricity effect and the HVA were more robust at 6 cpd than at 4 cpd. These findings are consistent with previous studies (eccentricity effect^5,59,60^; HVA^4,5,9,15,55^). A parallel pattern emerged for gain: the eccentricity effect and HVA were stronger at 6 cpd. This shared SF dependence reveals a qualitative link between performance and underlying computations, which likely reflect neural constraints: peripheral neurons prefer lower SFs in both humans^61,62^ and non-human primates^46,63,64^. Higher SFs place greater demands on spatial resolution, taxing locations with lower resolving power and thereby amplifying performance differences across eccentricity— and potentially around polar angle as well. The distinctions in underlying computations for eccentricity effect and polar angle asymmetries are consistent with a recent study, which using reverse correlation and a noisy-observer model, showed higher sensitivity to signal orientation and SFs and lower internal noise at the fovea than the parafovea^54^. In contrast, only orientation sensitivity varies around polar angle, mirroring performance^65^. The gain —an overall measure of how much the signal is amplified (current study) — and featural representation —the sensitivity profile to different features^54,65^— provide complementary descriptions of how the system enhances relevant input. Moreover, internal noise estimates obtained using the perceptual template model in the current study, and those derived from a noisy-observer model ^54,65^, quantify the same system-level property and together offer a more complete account of internal variability across the visual field.

These distinctions in underlying computations may reflect the interplay of four neurophysiological factors: (1) the number of neurons co-activated by visual stimuli, (2) the tuning properties of individual neurons (e.g., sensitivity and selectivity for visual features), (3) the intrinsic neural noise of individual neurons (e.g., firing variability and dendritic noise^47^), which contributes to system-level internal noise, and (4) correlations in activity across neurons^48,49^. By linking behavioral results to model-derived system-level computations, our results provide inferences about how these neurophysiological properties vary throughout the visual field.

Across eccentricity, the decrease in gain is consistent with two neurophysiological factors: First, as eccentricity increases, the cortical surface area stimulated by a given visual input decreases (e.g^26–31^), and thus, assuming uniform cortical density^66^, fewer neurons are recruited. Second, fewer neurons are tuned to high SFs at peripheral than central locations^46,61,63,64^ and receptive fields become larger^67^. Pooling from fewer and less well-tuned neurons likely result in weaker signal amplification and thus lower gain.

Whereas gain was substantially reduced from the fovea to the perifovea, it did not differ between the fovea and the parafovea. This result may seem surprising given the known decline in spatial resolution with eccentricity, but it may reflect an interaction between broader neural tuning and increased neural correlation. The increase in neural correlation at the parafovea can be attributed to a lower proportion of orientation-selective neurons^68^ and fewer parvocellular neurons^69^, which exhibit broader selectivity for SF^70^, than in the fovea. Together, these factors would offset the tuning advantage at the fovea, resulting in similar gain as at the parafovea. However, as eccentricity increases towards the perifovea, tuning continues to decrease while correlations remain high, resulting in a substantial drop in gain, consistent with our observations. The increase in internal noise with eccentricity may reflect a tradeoff between neural noise and neural correlation: The more correlated the neurons’ activity, the less likely their noise is to average out^48,49^. Although direct measurements of intrinsic noise across eccentricity are lacking, even if the noise of single neurons remains constant, increased neural correlation from fovea to periphery will result in greater system-level internal noise. This connection between higher neural correlation and higher internal noise is supported by findings that attention is associated with both lower neural correlations^71,72^ and reduced internal noise (e.g^40^).

The present study also provides the first assessment of nonlinearity across eccentricity. Here, nonlinearity —which reflects the system’s sensitivity to amplified signals (modeled by the power function in the perceptual template model)— exceeded 1 at all locations. Importantly, models without a nonlinear component, including the linear amplifier model^42^ and variants of perceptual template models^40,43^, failed to account for our data. These results underscore the need to consider nonlinear computations to explain visual performance. Further research is needed to determine whether this enhanced foveal nonlinearity shares neural mechanisms with the higher gain and lower additive noise observed at central locations.

Around polar angle, only gain varied and reflected both the HVA and VMA. Higher gain at better-performing locations may arise from greater cortical surface area^28–32,57,73^—implying more neurons, smaller population receptive field sizes^28,31^, and a preference for higher SFs^61,62,65^. This observed gain variation constrains hypotheses about how neural correlations vary around polar angle: robust gain variation around polar angle suggests that better-performing locations not only have superior tuning but also exhibit weaker neural correlations.

This interpretation on neural correlation, in turn, informs the observed invariance in internal noise around polar angle. As discussed in the eccentricity section, such invariance may reflect a tradeoff between neural correlation and neural noise. Thus, better-performing locations with weaker neural correlations —as suggested by our gain results—should have higher intrinsic neural noise. Although it may seem counterintuitive that neurons at better-performing locations could have higher neural noise, increased neural noise can be advantageous, e.g., it can facilitate the detection of subthreshold signals^74^. Future studies should map both neural correlations and neural noise across the visual field to characterize their interplay.

In sum, the observed gain variation across both eccentricity and polar angle likely results from an interplay of three factors at better-performing locations: a greater number of neurons, preference for higher SFs, and stronger neural correlations. In contrast, variation in internal noise across eccentricity —and its invariance around polar angle— may be jointly influenced by neural correlations and intrinsic neural noise, which our results suggest varying across the visual field. The relative impact of these neural factors remains an empirical question. A perceptual template model with population coding showed that neural correlations have a greater impact on performance than mere amplification or sharpening of tuning functions^75^.

Our findings may help explain why M-scaling eliminates eccentricity effects but not polar angle asymmetries in contrast sensitivity^7^. While M-scaling normalizes the cortical surface area stimulated, it may not equate other neurophysiological properties. For example, increasing stimulus size and thus the number of neurons reduces neural correlation^48,76^ and may diversify tuning profiles, further lowering such correlations. This reduction in neural correlation in the perifoveal region likely decreases system-level internal noise and therefore eliminates the eccentricity effect. In contrast, because internal noise appears uniform around polar angle, polar angle asymmetries—which are primarily driven by gain variations—may persist despite M-scaling.

It has been proposed that asymmetries between the upper and lower hemifields may relate to functional specialization, where the upper visual field primarily processes distant objects and the lower visual field processes near objects^77^. Polar angle asymmetries in contrast sensitivity^4–8^ and acuity^8,9^ decrease gradually with angular distance from the cardinal meridians, suggesting that whereas the near-versus-far specialization contributes to meridional asymmetries, it cannot fully account for the polar angle variations. We therefore hypothesize that gain-related polar angle asymmetries progressively diminish along non-cardinal meridians and disappear along the intercardinal meridians.

Previous studies have revealed ontogenetic and phylogenetic factors in these visual field asymmetries. Ontogenetically, children show an HVA both behaviorally^78^ and in cortical surface area^31^ but lack a behavioral^78^ or cortical^31^ VMA. The HVA progressively strengthens through adolescence, whereas the VMA emerges and becomes as pronounced as that of adults in late adolescence^79^. Phylogenetically, non-human primates show the typical HVA but an inverted VMA in visual performance. Evolutionary differences in locomotion, navigation, and tool use, which result in distinct visuomotor interactions at specific visual field locations, may contribute to shaping the organization of the visual field^80^.

In conclusion, by linking our novel behavioral data to underlying system-level computations, this study provides insights into the neurophysiological architecture of the visual field and generates novel, testable hypotheses. We found that visual field heterogeneities arise from distinct system-level computations. The eccentricity effect is driven by a combination of reduced gain and nonlinearity alongside increased additive noise, consistent with changes in cortical magnification and receptive field properties. Polar angle asymmetries are associated with differences in gain, but not in internal noise or nonlinearity. Together, these findings highlight a fundamental distinction in how the visual system implements computations across eccentricity versus and around polar angle, advancing our understanding of visual processing throughout the visual field.

## Materials and Methods

### Observers

Twelve observers (9 females; age mean=26.44; SD=3.24) participated in this experiment. All observers except two authors (S.X. and Q.C.) were naïve to the study’s purpose. All observers had normal or corrected-to-normal vision and provided written informed consent. The experimental methods complied with the Helsinki Declaration and were approved by the Institutional Review Board for Human observers at New York University.

### Apparatus

All experimental stimuli were generated using MATLAB 2020b (MathWorks) and the Psychophysics Toolbox^81^, and displayed on a gamma-calibrated CRT monitor (40 cm by 30 cm, 1,280 by 960 pixels, 100 Hz, 26 cd/m^-2^ background luminance). Each pixel extends 0.03 x 0.03 dva squared. The display was calibrated using a Photo Research (Chatsworth, CA) PR650 Spectracolorimeter to generate linear lookup tables for this experiment. The display resolution was 32 pixels per degree of visual angle. An EyeLink 1000 eye-tracker was positioned in front of observers to track gaze position. Observers were seated in a dark room with their head stabilized using a chin rest 57 cm from the display monitor.

### Stimuli

The stimulus presented in each trial consisted of a target, sandwiched between two streaks of dynamic noise patches. The target was a Gabor patch—a sinusoidal grating modulated by a Gaussian envelope—with luminance ( ) defined at each pixel position (and, relative to the patch’s center) (**Fig. 2A**, *inset*):

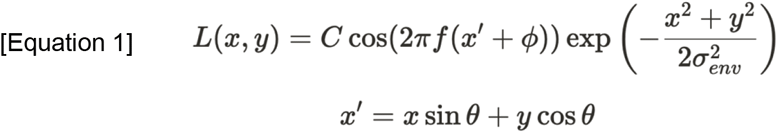

and range from −48 to 48 pixels (diameter=3 dva). represents linear contrast, either sampled from a staircase procedure or picked at a fixed value (Procedure). denotes SF (0.125 or 0.1875 cycles per pixel, equivalent to 4 and 6 cpd). denotes orientation in radians, either counterclockwise ( /4) or clockwise (−/4) with equal probability on each trial. denotes the phase which was randomized on each trial. specifies the Gaussian envelope size (=32 pixels, or 1 dva, one-third of the grating size).

The Gabor was embedded in dynamic noise, which were generated independently on each frame by sampling pixels from a Gaussian distribution with standard deviation ( ) and centered at 0 (hence, the RMS contrast of the noise patch equals) (**Fig. 2A**, *inset*):

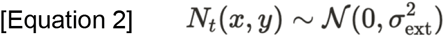

On each trial, could be one of nine possible values (0, 0.0275, 0.055, 0.0825, 0.11, 0.165, 0.22, 0.33, 0.44).

The possible stimulus locations were indicated by placeholder (**Fig. 2A**, *first display*): one at the fovea, and four along the horizontal (Left: 0°; right: 180°) and vertical meridians (upper: 90°; lower: 270°) at either 4° (parafovea) or 8° (perifovea) eccentricity. Each placeholder consisted of four corners (line length=0.75°).

### Procedure

**Fig. 2A** shows the trial sequence. Each trial began with all placeholders and a fixation dot (radius=0.1° of visual angle) at the center of a gray background. A streak of dynamic white noise was then presented at each location for a total duration of 630 ms. On each trial, a target Gabor was presented for 30 ms at one of the possible locations, 300 ms after the onset of the dynamic white noise, while other locations were only filled with noise. The streak of dynamic white noise persisted for 300 ms following target offset. The target location was marked by a boldened placeholder at the onset of the target Gabor, and until the observer’s response, eliminating location uncertainty.

Target location, orientation, and external noise levels were randomly selected on each trial. The Gabor SF (4 or 6 cpd) was constant within each experimental session. Observers performed an orientation discrimination task, pressing one of two keys to indicate whether the Gabor was tilted counterclockwise or clockwise (±45°). Task difficulty varied with stimulus parameters—Gabor visibility increased, hence tasks got easier, with higher contrast and lower noise standard deviation. Observers received auditory feedback after making a response: a high pitch for correct and a low pitch for incorrect responses. Their average accuracy was displayed at the end of each block.

We collected data to estimate contrast thresholds in two steps. First, at each noise level and location, we conducted a titration procedure using four interleaved staircases (two 1-up-2-down and two 1-up-3-down) starting from different contrast levels, with 40 trials per staircase. Second, based on Weibull fitting, we collected 30 to 40 additional trials at selected Gabor contrast levels— both within the dynamic range and at anchoring levels. We interleaved external noise and locations for all observers. Each observer completed 12,600 trials across approximately eight one-hour sessions for each SF. Nine observers were tested with both 4 cpd and 6 cpd stimuli, three were tested only at 6 cpd. They completed 25,200 trials in approximately 16 sessions. Stimuli generation and response recording were conducted with MATLAB. Data analysis and visualization were conducted in both MATLAB and R (R Core Team, 2024; RStudio v2024.12.1+563).

### Eye-tracking

We monitored the gaze position of the right eye in real-time during each trial using an SR Research Eyelink 1000 Desktop Mount eye tracker at a sampling rate of 1000 Hz. Stimulus presentation was contingent on maintaining central fixation from the onset until the offset of the dynamic white noise. Trials were terminated and re-queued to the end of the block if observers’ gaze deviated more than 1.5° of visual angle from the fixation center.

### Estimate contrast thresholds

For each noise level and location, we constructed psychometric functions by fitting a Weibull function to trials tested at varying Gabor contrast and corresponding binary responses (**Fig. 2B**). Trials collected through titration were binned into 10 bins based on percentile. Following titration, we collected additional trials at 2 to 3 fixed signal contrast levels per noise condition. These contrast levels were manually selected based on preliminary titration fits to improve the robustness of the psychometric function fitting.

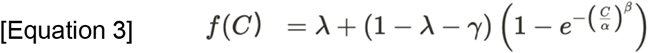

denotes accuracy as a function of Gabor contrast for each bin.,, and represent the function’s threshold at 75% accuracy, steepness, lapse rate, and chance performance, respectively. Model parameters were estimated using constrained nonlinear optimization via MATLAB’s fmincon() function. Parameter bounds were set as follows: thresh (\alpha) ranging from 1% to 100%, steepness ( ) from 0.01 to 100, guess rate ( ) from 0.45 to 0.55, and lapse rate (lambda) from 0 to 0.1, in 100 steps.

To balance the trade-off between trial count and fitting quality, we added “stabilizer” trials at low contrast levels (50 trials each at 0.01% and 0.1% Gabor contrast, enforcing 50% accuracy). At these low contrast levels, all observers were expected to perform at chance level, assuming no bias in reporting leftward or rightward orientation. The stabilizer trials were used only for fitting purposes and their trial counts were not included in the TvN fitting weights.

### Perceptual template model characterizes TvN curves

For each observer, we fitted a simplified version of the perceptual template model^40,41^ to contrast thresholds ( ) as a function of external noise levels ( ) at a given performance level ( ) (**Fig. 2C**):

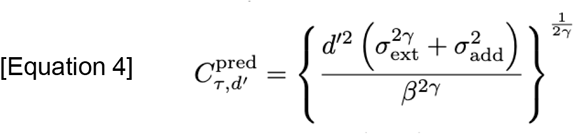

The model includes three key free parameters: (gain), a scaling factor reflecting how well the observer’s template matches the signal; (additive noise), the standard deviation of a Gaussian noise component independent of the stimulus strength; (nonlinearity), describing how the template-matched output is amplified. Note that the original full model also included a multiplicative noise parameter ( ) which reflects how internal noise scales with signal contrast^41^. By doing a model comparison we found that the reduced model excluding multiplicative noise outperformed the full model (p < 0.05, Bonferroni-corrected) (see Model Comparison below). Therefore, we proceeded with the more parsimonious three-parameter model (**Fig. 2C**).

#### Model Comparison

We evaluated the need to include all four parameters for the perceptual template model by conducting a nested model comparison between the full model and 15 reduced models, each systematically excluding one or more parameters. The full model devised and tested by Lu and Dosher is:

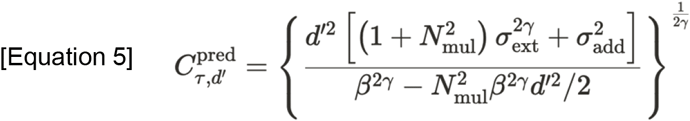

When excluded from fitting, multiplicative ( ) and additive ( ) internal noise were fixed at 0, and gain ( ) and nonlinearity ( ) were fixed at 1. We fitted each candidate model to the seven locations we tested (fovea, horizontal meridian, upper and lower vertical meridian at 4° and 8° eccentricity). We first excluded models that failed to account for the data (indicated by negative R-squared), which occurred when gain and/or additive noise were omitted. For the remaining four models that included both gain and additive noise, we quantified their goodness of fit by the Bayesian Information Criterion (BIC):

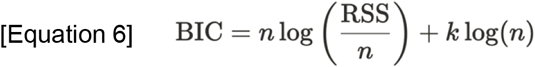

RSS is the sum of squared residuals specified above, is the number of data, and is the number of parameters. Lower BIC scores indicate better model fits while penalizing overfitting. We fitted the full set of perceptual template models to each location group independently for each observer, allowing all parameters to vary freely across locations within each location group. Delta BIC was calculated by subtracting the minimum BIC score from each candidate model’s BIC score. Finally, we fitted linear mixed-effects models to delta BIC values to evaluate the fixed effect of model type with random effects specified across observers:

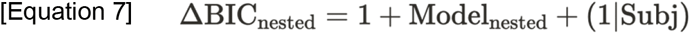

The reduced model excluding multiplicative noise outperformed the full model (p < 0.05, Bonferroni-corrected). Therefore, we proceeded with the more parsimonious three-parameter model.

When fitting the perceptual template model, additive noise was calculated in log10 units because they are in contrast units. Model fitting was implemented by MATLAB’s fmincon() function, which minimized the sum of squared residuals (RSS) between the measured and predicted thresholds. Prior to squaring, residuals were calculated on log contrast values and weighted by the number of trials per noise level ( ).

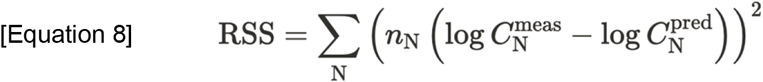

Parameter bounds were decided according to the literature^9,41^: multiplicative noise ( ) was constrained between 0 to (to ensure a positive denominator in equation (4)), additive noise ( ) ranged from 10^−8 to 1 in log steps, gain ( ) was bounded between 0 to 5, and nonlinearity ( ) ranged from 0 to 5.

To constrain model estimation^40,41^, we obtained thresholds ( ) at three performance levels =0.77, 1.05, 1.35, where, corresponding to 65%, 70% and 75% accuracy ( ), respectively. Only thresholds at 75% correct were shown as TvN functions (**Fig. 3**). This approach was originally intended to constrain the model by ruling out substantial reductions in internal multiplicative noise as a plausible mechanism of performance variation^43^. Although we ultimately adopted a simpler version of the model that does not require this constraint anymore, we retained the method for consistency with prior literature.

To examine visual field heterogeneity (eccentricity effect, HVA, and VMA), we grouped locations according to the effect of interest (see Statistical Analysis-Location groups below), then fitted perceptual template models to all locations within each location group.

### Statistical Analysis

#### Location groups

At each noise level, we derived thresholds for eccentricity and meridian by combining data across locations. To represent the horizontal meridian at each eccentricity, we collapsed trials from the left and right locations and fit a single psychometric function. Thresholds for the vertical meridian were computed as the mean of the upper and lower vertical meridian. Thresholds for each eccentricity were computed as the means across the horizontal and vertical meridian.

We grouped visual field locations to systematically examine the eccentricity effect, HVA, and VMA. To assess the eccentricity effect, we compared thresholds at the fovea, 4°, and 8° along the horizontal and vertical meridians separately (i.e., three locations in each group), allowing us to examine whether the meridian modulates eccentricity-related changes. To evaluate polar angle asymmetries, we compared thresholds between the horizontal and vertical meridians (for HVA) and between the lower and upper vertical meridians (for VMA) at 4° and 8° eccentricities, with four locations included in each group. In addition, we defined four supplementary groups to isolate each asymmetry at individual eccentricities, enabling follow-up analyses when needed.

Within each group, we fit the perceptual template model to thresholds across external noise levels to estimate the underlying system-level computations. Grouping was essential to isolate location-specific computations without conflating repeated measures. Some locations appeared in multiple groups (e.g., the fovea appeared in both eccentricity groups), leading to multiple parameter estimates per location. One-way ANOVAs confirmed that these estimates did not differ across groups (all *p*s > 0.1), and all were retained for further analysis.

#### Linear mixed-effects model

We used linear mixed-effects models for three sets of analyses: (1) verifying the eccentricity effect and polar angle asymmetries, (2) determining the optimal parameter configuration of the perceptual template model, and (3) analyzing the spatial pattern of computations. This approach was chosen for two reasons. First, linear mixed-effects models handle unbalanced datasets effectively, which was optimal given our mixed design: a between-observer factor (nine observers were tested at both 4 and 6 cpd, and an addition of three people were tested only at 6 cpd) and a within-observer factor (three thresholds were derived per location x noise per observer). Second, linear mixed-effects models account for random effects by including observer-specific intercepts, capturing individual variability across conditions.

All analyses were conducted in RStudio (v2024.12.1+563) using R. Linear mixed-effects models were fitted using the lmer() function from the lmerTest package, with maximum likelihood estimation and degrees of freedom calculated using the Satterthwaite approximation, which improves accuracy under unequal sample sizes and the inclusion of random effects. We used the anova() function to perform Type III ANOVA tests for assessing main effects and interactions, and the emmeans package for post-hoc comparisons with Bonferroni corrections. Model fit was evaluated using conditional R-squared values, which reflect both fixed and random effects.

#### Visual field heterogeneity for performance

Before examining the computations, we first verified that eccentricity effects and polar angle asymmetries were robust for contrast threshold in the absence of external noise, consistent with previous studies. We defined “performance” as the intercept threshold—the threshold measured without external noise— the standard metric in studies without external noise. To align with conventional performance definitions, where higher values indicate better performance, we took the absolute value of log-transformed thresholds. We assessed how performance varied across visual field locations within each location group and whether SF modulated these variations. To verify eccentricity effects, we separately modeled performance within each location group using linear mixed-effects models, with fixed effects for eccentricity, SF, and their interaction:

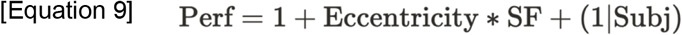

To verify HVA and VMA, we fit separate linear mixed-effects models for each asymmetry. For HVA, we tested the fixed effects of *meridian* (horizontal vs. Vertical meridian), eccentricity (4° and 8°) and SF and their interactions. For VMA, we modeled the fixed effects of *location* (lower vs. upper vertical meridian), eccentricity and SF, and interactions among these factors.

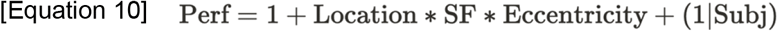

#### Variation and contribution

We investigated how these parameters underlie visual field heterogeneity throughout the visual field (eccentricity effect, HVA, and VMA) by examining the variation and contribution of neural computations.

First, to analyze how each computation varies across locations and how these variations interact with SF, we fitted a linear mixed-effects model to each parameter. To assess variation across eccentricity, similar to equation (6), the model assesses the interaction between eccentricity (4° and 8°) and SF (4 and 6 cpd):

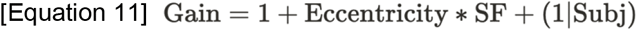

To assess variation around polar angle, similar to equation (7), the model assesses the interaction among location (meridian for HVA and location for VMA), SF and eccentricity:

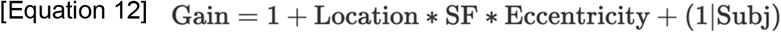

Both models had random intercepts for observers. Regression slopes were constrained to be constant, enabling assessment of overall trends.

Then, we examined how each parameter modulates performance. We analyzed the fixed effect of each parameter. Using gain as an example:

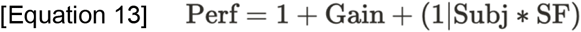

This model includes a random intercept for each location-SF combination. Hence, any observed fixed effect cannot be attributed to differences across locations or SFs. We chose not to include random slopes because this analysis aims to estimate the overall fixed effect across locations and SF. The model was fitted to the seven locations that cover the isoeccentric locations at the tested eccentricity range.

After determining whether each parameter enhanced or impaired performance, we assessed the independent contribution of each parameter to each performance heterogeneity. First, we quantified the extent of asymmetry for both performance and parameters using a location difference index (LDI)—the arithmetic difference between two locations divided by their sum. This index provides a normalized measure of location variation. By design, the first of the two values in the LDI calculation was always taken from the location with higher performance. For example, when comparing the fovea and 8°, both the numerator and denominator of the LDI were computed as the difference between fovea and 8°. We then fitted linear mixed-effects models to LDI values of each parameter (e.g., gain) and that of performance, and the resulting fixed effects indicate contribution of each parameter.

For gain and additive noise, we computed one-tailed empirical *p*-values (see Bootstrapping section for definition) because the directionality of their contribution to performance heterogeneity could be inferred from the directionality of their modulation on performance. Specifically, if better-performing locations have higher gain, then gain is assumed to enhance performance. In this case, both the performance LDI and gain LDI would be positive, and a stronger difference in performance would correspond to a larger difference in gain—resulting in a positive correlation. Conversely, if a parameter impairs performance (e.g., additive noise), a negative correlation would be expected. If a parameter does not modulate performance (e.g., nonlinearity), we evaluated the two-tailed empirical *p*-value.

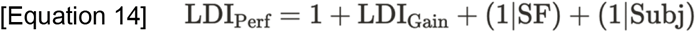

#### Bootstrapping

To derive confidence intervals and assess statistical significance, we performed bootstrapping beginning at the level of threshold estimation. In each iteration, we resampled trials with replacement at each location and noise level separately. Psychometric functions were then fitted to the resampled data to estimate contrast thresholds at each noise level. These thresholds were subsequently fitted with the perceptual template model within each location group to estimate the three model parameters. Finally, statistical analyses were performed across observers to assess modulation, variation, and contribution effects. Both the resampling and the two-stage model fitting were conducted independently for each observer. This procedure was repeated 1,000 times. We reported the observed statistics for all tests (*F*-values, *t*-values, effect sizes, slopes) and *p*-values (*p*_*obs*_) from the non-bootstrapped data, along with 95% confidence intervals (excluding *p*-values) from the bootstrap distribution. For pairwise comparisons and correlation analyses, empirical two-tailed *p*-values (*p*_*emp*_) were computed as twice the smaller proportion of bootstrapped differences (and slopes) greater or less than zero, and empirical Cohen’s *d* was computed as the median of the bootstrapped differences divided by the standard deviation of those differences. For correlation analyses assessing contribution effects, the choice between one-tailed and two-tailed empirical *p*-values depended on the parameter’s modulation effect: if the parameter significantly modulated performance in a specific direction, a one-tailed *p*-value was used; otherwise, a two-tailed *p*-value was applied.

## Acknowledgments

We thank David Tu and Zhong-Lin Lu for their helpful comments. This research is funded by NIH R01-EY027401 to M.C., Vision Training Grant 5T32EY007136-30 to NYU, and NIH F31-EY036732 to S.X.

## Author Contributions

S.X., A.B., J.A. and M.C. conceived and designed the study. S.X. , A.B. and J.A. wrote the experimental code. J.A. collected preliminary data. S.X., A.B. and Q.C. collected and analyzed the experimental data. S.X. drafted the manuscript. M.C. administered and supervised the project; provided extensive editorial input and critical revisions throughout the writing process. All authors provided feedback, contributed to editing and approved the final version of the manuscript.

